# Designed Variants of ACE2-Fc that Decouple Anti-SARS-CoV-2 Activities from Unwanted Cardiovascular Effects

**DOI:** 10.1101/2020.08.13.248351

**Authors:** Pan Liu, Xinfang Xie, Li Gao, Jing Jin

**Author notes:** Correspondence to: Jing Jin, 303 E. Superior Street, Room 8-518, Chicago, IL 60611.

## Abstract

Angiotensin-converting enzyme 2 (ACE2) is the entry receptor for SARS-CoV-2, and recombinant ACE2 decoys are being evaluated as new antiviral therapies. We designed and tested an antibody-like ACE2-Fc fusion protein, which has the benefit of long pharmacological half-life and the potential to facilitate immune clearance of the virus. Out of a concern that the intrinsic catalytic activity of ACE2 may unintentionally alter the balance of its hormonal substrates and cause adverse cardiovascular effects in treatment, we performed a mutagenesis screening for inactivating the enzyme. Three mutants, R273A, H378A and E402A, completely lost their enzymatic activity for either surrogate or physiological substrates. All of them remained capable of binding SARS-CoV-2 and could suppress the transduction of a pseudotyped virus in cell culture. This study established new ACE2-Fc candidates as antiviral treatment for SARS-CoV-2 without potentially harmful side effects from ACE2’s catalytic actions toward its vasoactive substrates.

## 1. Introduction

As COVID-19 pandemic is still unfolding and no specific antiviral treatments are available, there is an unmet need to explore new drug candidates that are effective and safe, and broad spectrum against the evolving virus. In addition to the ongoing clinical trials of repurposed compounds and patient-derived antibodies, new drugs are being developed through targeted screening and rational design.

One of the focuses is on drug candidates that target receptor-mediated viral entry. A diverse group of human coronaviruses including SARS-CoV of 2002, HCoV-NL63 of 2004 and SARS-CoV-2 of COVID-19 rely on their spike proteins to bind ACE2 cell receptor as the first step in viral entry[1-4]. It has been shown that soluble ACE2 at generous abundance as compared to viral concentration can lower infectivity of cultured human cells, similar to experimental anti-spike antibodies[5-7]. Prior to the pandemic, one of the human recombinant soluble ACE2 (hrsACE2) drugs developed by Apeiron Biologics and GlaxoSmithKline (GSK) had completed Phase I and Phase II clinical trials for human pulmonary arterial hypertension and acute respiratory distress syndrome (ARDS) [8, 9], and is now repositioned for investigational treatment of COVID-19 (ClinicalTrials.gov identifier: NCT04335136).

The current focus has been on improving binding affinity, pharmacokinetic/pharmacodynamic (PK/PD), antiviral specificity and neutralization efficacy of ACE2-based biologics through bioengineering design[6, 10-12]. Also in a different context unrelated to COVID-19, our group had previously constructed a chimeric fusion between the ectodomain of ACE2 and the Fc segment of IgG1 (“hinge” plus CH2 and CH3 regions) (**Fig1A**). In keeping with a well-recognized function of Fc to extend protein half-life through its cognate neonatal Fc receptor (FcRn), ACE2-Fc has improved pharmacokinetics as compared to untagged soluble ACE2[13]. The enzymatic activity of ACE2 in the fusion degraded angiotensin II (AngII) and rendered blood pressure control for up to two weeks. In comparison to its untagged counterpart to treat COVID-19, ACE2-Fc is predicted to offer superior pharmacological benefits, which make it also suitable for prophylactic usages by frontline healthcare workers and caregivers[11].

**Figure 1.**
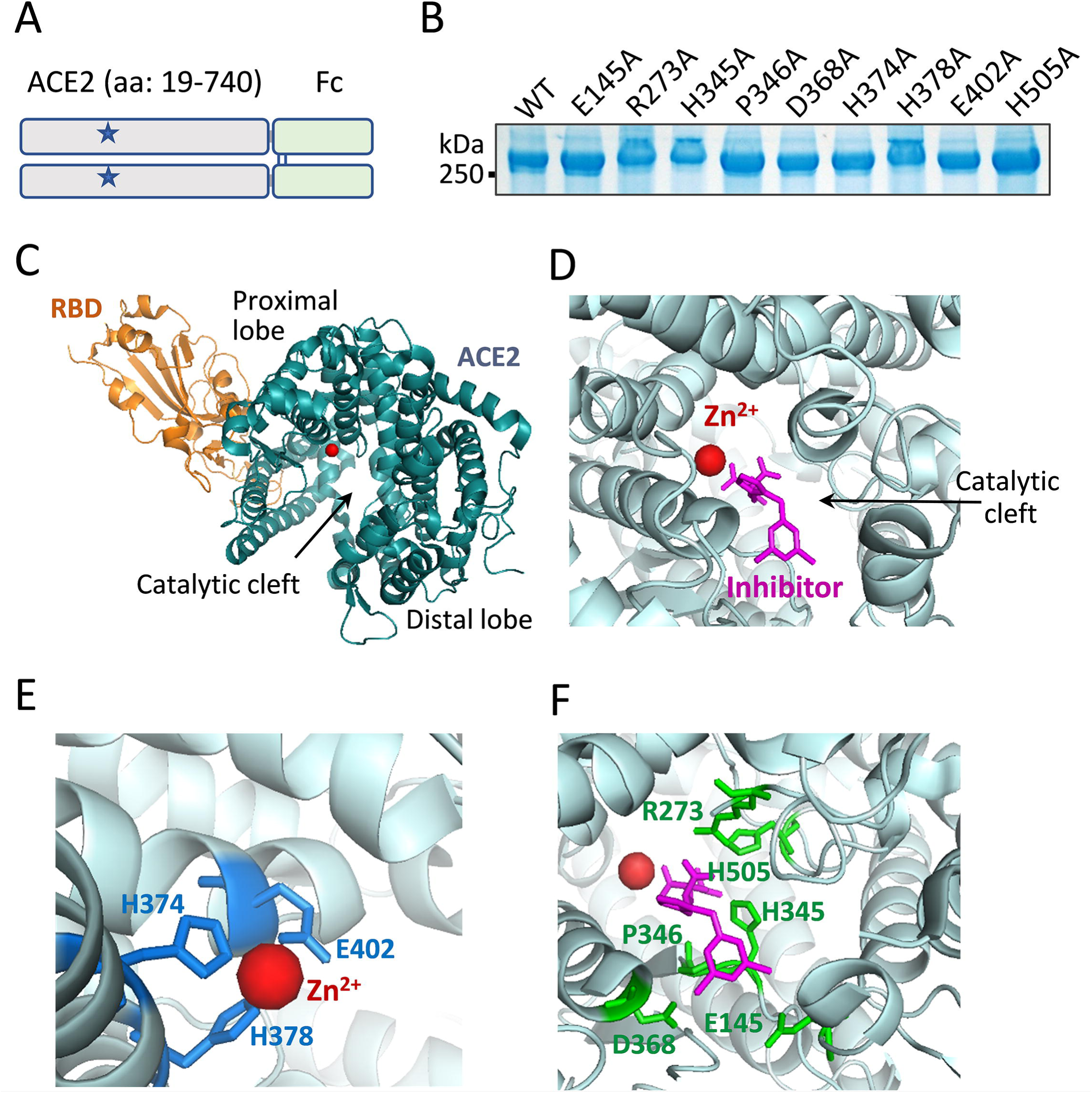
Mutagenesis strategies for catalytic inactivation of ACE2. **A**. Schematic representation of chimeric ACE2-Fc. The ectodomain sequence from amino acid 19 to 740 of human ACE2 is fused to the N-terminus of Fc domain of human IgG1, which forms a dimer via disulfide bridges (blue lines). The overall plan was to make individual point mutations (denoted as stars) of the catalytic site to inactivate the ACE2 peptidase activity. **B**. Mutant ACE2-Fc proteins, each carries a single alanine substitution of a selected residue, were produced by HEK-293 cells. As predicted, the proteins ran above the 250 kDa marker under nonreducing condition. **C**. The cocrystal structure (PDB: 6M0J) shows SARS-CoV-2 spike RBD binding of ACE2. ACE2 exists in a clam shell-like configuration holding a catalytic cleft between its proximal and distal lobes. A zinc ion resides within the proximal lobe of the cleft void. **D**. In an inhibitor (MLN-4760)-bound structure of ACE2 (PBD: 1R4L), the inhibitor induced a conformational change of the catalytic cleft to adapt a ‘closed’ configuration[40]. **E**. Three proximal lobe residues H374, H378 and E402 formed interactions with the zinc ion. **F**. The side chains of six proximal and distal lobe residues, E145, R273, H345, P346, D368 and H505 (in green) formed direct interactions with inhibitor MLN-4760 (in magentas).

Our study attempts to address another potential drawback of hrsACE2 biologic. Although it was originally believed that the catalytic activity of ACE2 delivered in an excess quantity through therapeutic hrsACE2 may alleviate ARDS based on mouse studies[8, 14], the relevance in human disease remains unclear. COVID-19 mortality is prevalent among patients with underlying conditions such as cardiovascular disease, diabetes and chronic lung disease [15-18], a large number of COVID-19 patients are on existing ACEI/ARB blockade therapy for preexisting cardiovascular and diabetic comorbidities[19]. There have been different opinions on whether RAAS blockade medications had improved or worsened COVID-19 recovery[20-24]. However, studies discovered a correlation between RAAS blockade and upregulation of endogenous ACE2 expression, causing concern of increased risk of SARS-CoV-2 infection[25-29]. The carboxypeptidase activity of therapeutic hrsACE2 hydrolyses a broad range of vasoactive hormonal substrates including AngII, apelin-13, bradykinin, among others, and exerts systemic RAAS blockade that affects the heart, the blood vessels, the kidney and the lung. Severe cases of SARS-CoV-2 infection frequently have multiorgan involvement [30-34]. In our view, the dual functions of investigational drug hrsACE2 to simultaneously act on viral neutralization and RAAS can potentially complicate clinical assessment of therapeutic efficacy. In order to achieve an exclusive antiviral function in an ACE2-derived biologic, we sought to modify ACE2 catalytic center to limit the catalysis of its vasoactive substrates.

## 2. Materials and Methods

### 2.1. Construction of ACE2-Fc mutant plasmids

DNA sequence encoding the ectodomain of human ACE2 (aa 1-740) was cloned from a human kidney cDNA library. DNA encoding human Fc of IgG1 has been described previously[13]. An in-frame fusion between ACE2 and Fc was constructed in pcDNA3 vector (Invitrogen, Carlsbad, CA). Site-directed mutagenesis by PCR was performed to create the panel of ACE2-Fc mutants. All mutants were confirmed by sequencing.

### 2.2. Recombinant protein expression and purification

The workflow for generating recombinant ACE2-Fc variants was similar to what had been reported before[13]. Briefly, HEK293 cells were transfected with individual ACE2-Fc variants by standard polyethylenimine (PEI) method. In transgene transfection studies, on day two of transfection cells were switched to serum-free DMEM. On day four, the culture media were harvested by centrifugation, and further concentrated using Amico Ultra Filters (Millipore, Billerica, MA). ACE2-Fc proteins were then purified by size-exclusion chromatography (SEC) using Superdex 200 Increase column (GE healthcare, Chicago, IL) and stored at −80°C until used in experiments.

ΔACE2-Fc Arg273Ala, His378Ala and Glu402Ala, and wild-type ACE2-Fc selected for scaled production were produced from clonal stable-expressing cells. The general method was described previously[13]. Following plasmid transfection of HEK293 cells, the cells were selected under 1 mg/mL G418 (Thermo Fisher Scientific, Waltham, MA) for ∼14 days until isolated cell colonies appeared in the dishes. Individual cell clones were seeded into 96-well plates. When cells reached 50-100% density in the wells, the culture media were tested for their ACE2-Fc contents using a custom ELISA (anti-ACE2 antibody [Abcam, Cambridge, MA] for capturing and anti-human IgG-Fc-HRP [SouthernBiotech, Birmingham, AL] for detection). Clones with the highest expression of recombinant ACE2-Fc variants were selected, and individually expanded to five 150 mm dishes. Once reached ∼90% confluency, the cultures were switched to serum-free DMEM medium. After 4-5 days the culture media were harvested, and further concentrated using a VIVAFLOW 200 filtration system (100,000 MWCO by Sartorius, Stonehouse, UK). Recombinant ACE2-Fc proteins were purified using SEC as described above.

### 2.3. ACE2 peptidase activity measured using surrogate Mca-APK(Dnp)

ACE2-Fc peptidase activity assay using surrogate fluorogenic substrate Mca-APK(Dnp) (Enzo Life Sciences, Farmingdale, NY) was performed in black microtiter plates. The reaction buffer contained 50⍰mM 4-morpholineethanesulfonic acid, pH⍰ =⍰6.5, 300⍰mM NaCl, 10⍰μM ZnCl_2_, 0.01% Triton X-100 and 20⍰μM of Mca-APK(Dnp). The total reaction volume was 100⍰μL at room temperature and the duration of the reactions were 20 min. Peptidase activities were calculated as fluorescence intensity at 320 nm excitation and 420 nm emission wavelength.

### 2.4. ACE2 peptidase activity measured using physiological substrates of AngII and apelin-13

We described the method previously and referred to the workflow as Phenylalanine Assay, which was tested using AngII and apelin-13 as substrates in reactions with ACE2[35]. It involves two coupled reactions, hydrolysis of the C-terminus phenylalanine residues from the substrates by ACE2 catalysis and the measurement of free amino acid phenylalanine using yeast enzyme of phenylalanine ammonia lyase (PAL) in a colorimetric assay (supplementary Figure S2).

The reactions were carried out using indicated concentrations of ACE2-Fc proteins together with either AngII or apelin-13 at indicated concentrations in reaction buffer containing 20 mM Tris-HCl, pH⍰= ⍰7.4, 136⍰mM NaCl and 10⍰μM ZnCl_2_. The first reaction was proceeded at 37°C after 20 min before stopped by 80°C heat inactivation for 5 min. The second reaction used a phenylalanine detection kit (Sigma-Aldrich, St. Louis, MO). 1 μL enzyme mix and 1⍰μL developer from the kit were added to the above reaction, which was allowed to proceed for 20 min at room temperature. Fluorescence intensity was measured at 5351nm excitation and 5851nm emission wavelength, and all reactions were performed in triplicate.

### 2.5. SARS-CoV-2 RBD binding assay

Recombinant viral RBD protein was purchased from Sino Biological (Beijing, China). ELISA wells were precoated with either PBS as controls or 100 ng/well of RBD protein. Serial concentrations of individual ACE2-Fc variants were added to the wells. After overnight incubation at 4°C, the wells were washed three times with TBST buffer before HRP-conjugated anti-human IgG-Fc secondary antibody was added. HRP reactions were developed with TMB substrate and the binding strength derived from OD450 (nm) readings of the reactions. The EC50 values were determined by log(agonist) vs. response nonlinear regression fit analysis (GraphPad Prism).

### 2.6. Inhibition of viral transduction with purified ΔACE2-Fc mutants

Spike (SARS-CoV-2) pseudotyped lentivirus with luciferase reporter gene was purchased from BPS Bioscience (San Diego, CA). The virus was used to transduce HEK293 with stable expression of receptor full-length ACE2. The stable cell line was created from plasmid expressing full-length human ACE2 in pcDNA3 vector. Followed a similar procedure for generating ACE2-Fc clones, stable HEK293 clones with receptor ACE2 expression was identified using immune detection of ACE2 in the cells. The ACE2-HEK293 cells were seeded at a density of 10,000 cells per well into white 96-well cell culture microplate one day before transduction. To test inhibition of viral transduction, 5 μL pseudotyped lentivirus were preincubated with 5 μL vehicle or serially diluted ACE2-Fc variants at 37 °C for 1 h and then added into the cells. After overnight incubation, the cells were refed with fresh medium and incubated for another 36 hours. Luciferase activity was measured using ONE-Glo™ Luciferase Assay System according to the manufacture’s protocol (Promega, Madison, WI). The IC50 values were determined by log(inhibitor) vs. response nonlinear regression fit analysis (GraphPad Prism).

### 2.7. ACE2-Fc pharmacokinetics in mice

Institutional Animal Care and Use Committee of the Northwestern University approved the animal procedure in this study (approved protocol number IS00009990).The general method for pharmacokinetic measurements were described previously[13]. Briefly, a bolus intravenous injection of ACE2-Fc proteins (0.5mg/kg body weight) was performed in 10 weeks old female BALB/c mice. Subsequently, serial blood samples were collected from tail bleeding at indicated time points. Collected blood samples were left undisturbed on ice, and sera were isolated by centrifugation at 6000 x g for 10 minutes at 4°C. The levels of ACE2-Fc in the sera were measured by ELISA using anti-ACE2 capturing antibody and anti-human IgG-Fc-HRP antibody for detection as described above.

## 3. Results

### 3.1. ACE2-Fc mutagenesis strategy to remove catalytic activity

We constructed an ACE2-Fc template by using the ectodomain of ACE2 fused with an Fc sequence (**Fig1A**). The chimeric fusion naturally formed a dimer of >250 kDa as expected (**Fig1B**). There is an extensive amount of information with regard to the structural characteristics of ACE2 in relationship to SARS-CoV-2 receptor-binding domain (RBD)[1, 2, 36-39]. SARS-CoV-2 binds a surface segment of ACE2 through the apex of its spike protein (**Fig1C**). ACE2 is a metallopeptidase that requires divalent cation such as zinc for activity. A Zn^2+^ ion is buried deep in the catalytic cleft within the proximal lobe relative to the viral binding site. Based on an inhibitor-bound structure of ACE2[40], both proximal and distal residues that line the catalytic cleft form interactions with the inhibitor, which occupies the presumed substrate pocket (**Fig1D**). Zn^2+^ is coordinated by three residues, His374, His378 and Glu402, which are the obvious choices for mutagenesis when making enzymatically inactive mutants[6] (**Fig1E**). In addition to these Zn^2+^-binding sites, we sought to look for catalytic residues in contact with the substrates that are further away from RBD binding segment. The inhibitor-bound structure indicates six residues that extend their side chains toward the substrate direction. These are Glu145, Arg273, His345, Pro346, Asp368 and His505 (**Fig1F**).

### 3.2. Substrate-dependent inactivation among ACE2-Fc mutants

Next, we made alanine-substitution of each residue including the three that bind Zn^2+^ and the additional six that potentially bind substrates (supplementary figure S1). Wild-type and 9 mutants of ACE2-Fc were produced using a HEK293 expression system as soluble proteins (**Fig1B**). Purified proteins were subjected to a set of enzymatic assays using a surrogate Mca-APK(Dnp) fluorogenic substrate and physiological substrates such as AngII and apelin-13 (**Fig2**). It should be noted that although the surrogate substrate Mca-APK(Dnp) is traditionally used for measuring ACE2 activity, the sequence does not resemble those of the physiological substrates, which share a Pro-Phe motif at their C-termini (Supplementary figure S2). Instead, the catalysis of AngII and apelin-13 by ACE2-Fc variants was measured by the hydrolysis rate of their C-terminus Phe amino acid[35].

**Figure 2.**
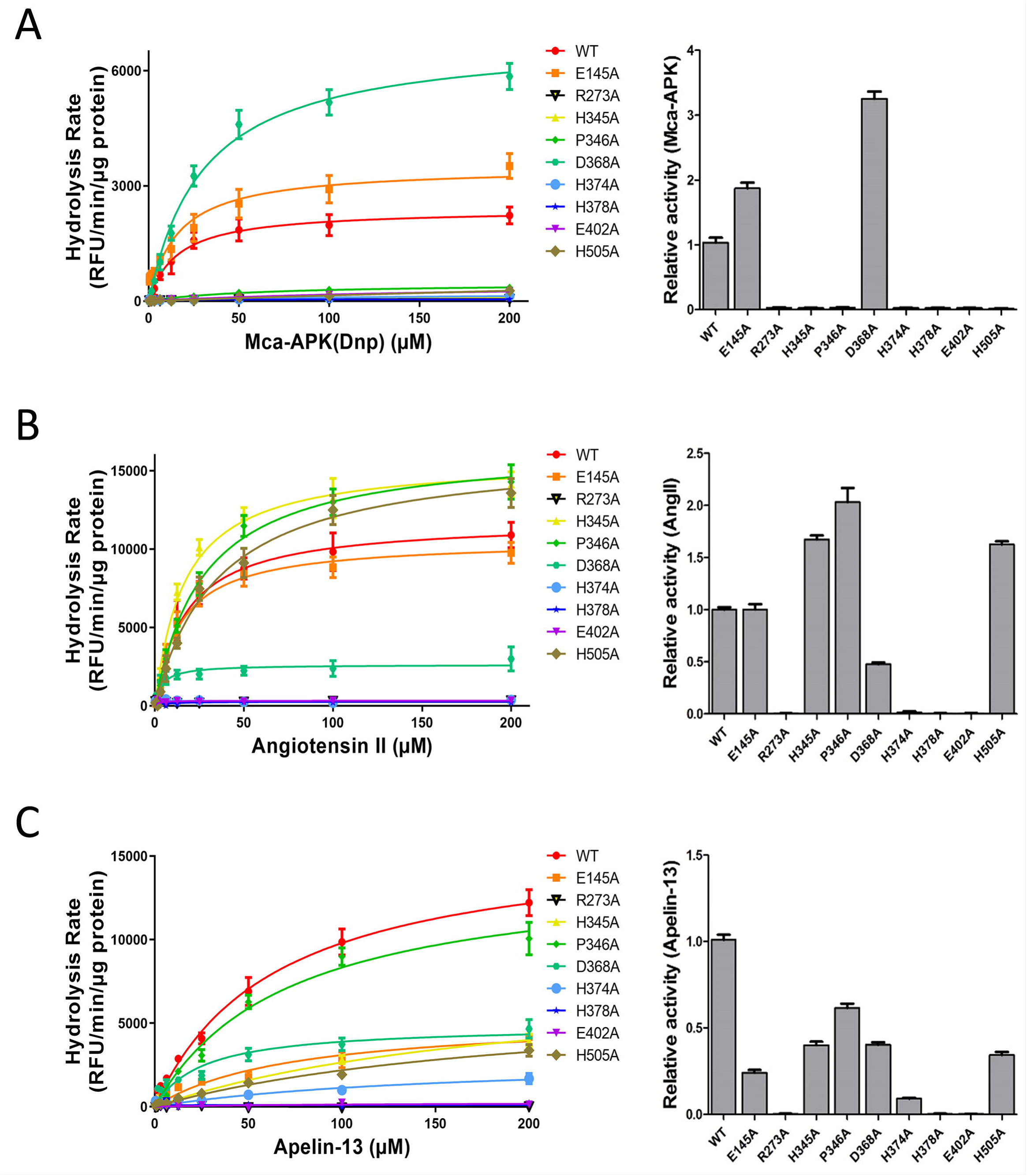
Substrate-dependent inactivation of ACE2-Fc peptidase among ACE2-Fc mutants. Three peptide substrates were tested in catalytic reactions with ten individual variants of ACE2-Fc (wild-type and 9 mutants). The reactions were carried out in two different ways. Left panels: the reactions were performed using a high amount of purified ACE2-Fc enzyme (100 ng) with varying concentrations of the substrates between 0.39 µM and 200 µM (x-axis). Right panels: a lower dose of 10 ng ACE2-Fc was incubated with a fixed amount of 2 nmole of Mca-APK(Dnp) or 10 nmole of AngII/Apelin-13. Reactions proceeded for a standard length of time of 20 min. **A**. Surrogate fluorogenic substrate Mca-AP↓K(Dnp) was tested (↓: cleavage site). ACE2-Fc peptidase activities were compared between wild-type and mutants. Seven out of the nine mutants showed a completely loss-of-activity. **B**. When AngII was used in reactions with the ACE-2-Fc panel, DRVYIHP↓F of AngII sequence was cleaved by ACE2, releasing a Phe/F. The assay detected the generation of amino acid Phe/F as the results of ACE2-Fc activity. **C**. Similar to AngII, Apelin-13 was cleaved by ACE2 between the proline(P)-phenylalanine (F) bond in its sequence, Pyr-RPRLSHKGPMP↓F. The rates of Phe/F release from the reactions were detected. Data are shown as the mean ± SD from triplicate experiments.

We performed Mca-APK(Dnp) measurements, alongside phenylalanine hydrolysis assays using AngII and apelin-13 as substrates to determine the enzymatic activities of the ACE2-Fc variants (**Fig2A-C**). In order to better characterize the catalytic performances of individual ACE2-Fc mutants relative to their wild-type counterpart, we conducted two types of enzymatic assays. The first method used an excess amount of the enzyme (100 ng) in reactions with varying concentrations of the three substrates. This would potentially detect low partial activity of enzymes. The second method had a lower amount of purified ACE2-Fc variants (10 ng each) to react with an excess quantity of the substrates (2 or 10 nmol: see Methods) in order to distinguish among mutants with high activities. As it turned out, the results from these two methods were to an extend in agreement with each other (**Fig2**: left panels compared to right panels). One of the surprising findings was that there was a clear evidence of substrate-dependent inactivation of individual mutations, particularly among those lining the inhibitor/substrate space. For instance, while Mca-APK(Dnp) showed no activity of 7 ACE2-Fc mutants, including 4 substrate-binding residues of Arg273Ala, His345Ala, Pro346Ala and His505Ala (**Fig2A**), His345Ala, Pro346Ala and His505Ala remained active toward AngII and apelin-13 (**Fig2B,2C**). In addition, His374Ala, one of the Zn^2+^-binding residues, retained a low level of activity against Apelin-13 (**Fig2C**). When all three substrates are considered, Arg273Ala, His378Ala and Glu402Ala were completely lack of peptidase activity. We referred these three mutants to as ΔACE2-Fc (Δ: loss-of-activity) and considered them as candidate variants emerged from the screen.

### 3.3. Binding affinities of individual ACE2-Fc mutants to SARS-CoV-2 receptor-binding domain

Next, we performed binding assays using individual variants against purified spike RBD protein. As expected, all mutants displayed similar levels of binding to viral RBD, considering the relatively minor changes from the point mutations made to ACE2-Fc (**Fig3**). Overall, the wild-type protein showed the highest binding affinity, whereas all three Zn^2+^-binding site mutants, His374Ala, His378Ala and Glu402Ala, had the lowest affinities to RBD. This is consistent with the expectation that the ion pocket is in proximity to the viral binding site on ACE2 (**Fig1C**), and also that changing ion-binding can potentially induce structurally instability of the protein. In contrast, Arg273Ala mutant of the substrate-binding pocket, which showed complete loss-of-activity towards all three substrates, is situated on the distal lobe and is less likely to affect desired viral binding.

**Figure 3.**
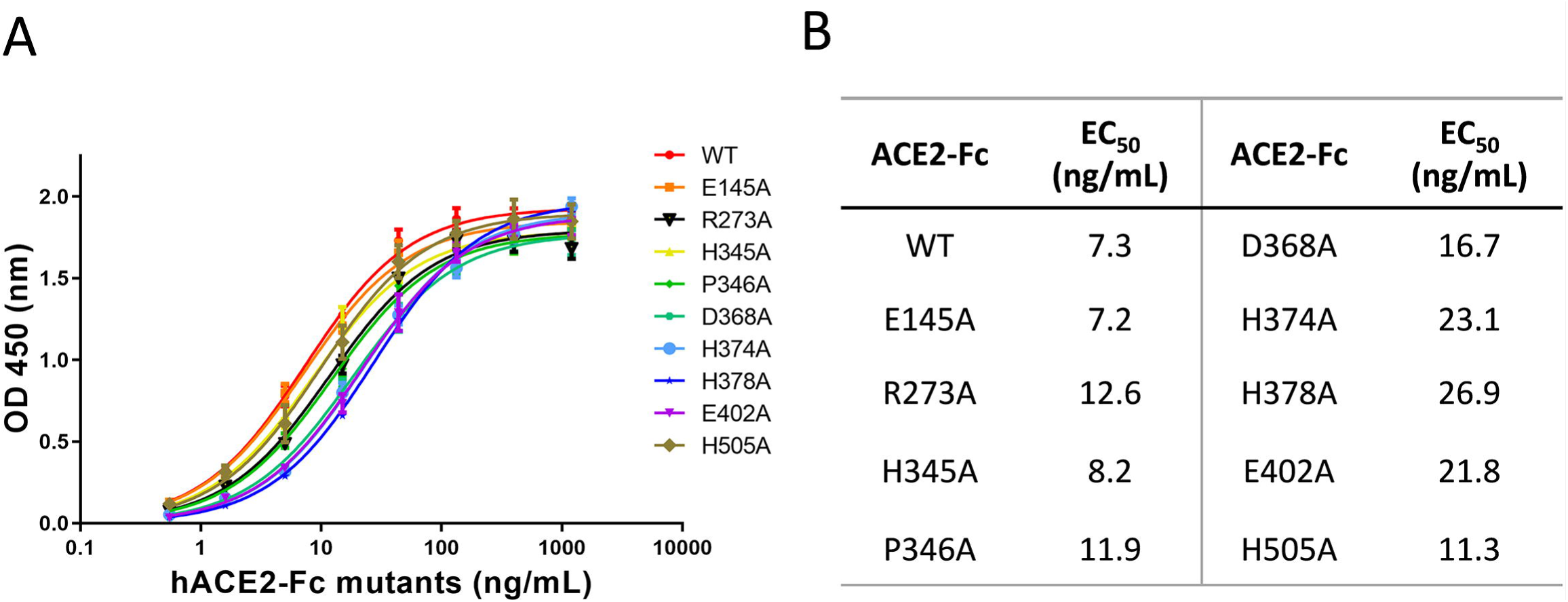
Binding affinities of individual ACE2-Fc variants to RBD of SARS-CoV-2 spike protein. **A**. A fixed amount of recombinant viral RBD protein was coated to an ELISA plate (buffer coated well were used as controls). Wild-type and nine mutants of ACE2-Fc were added to the wells at varying concentrations between 0.5 ng/mL and 1200 ng/mL (x-axis). Binding was determined by the difference in signal intensity between the RBD-coated and the corresponding control wells. **B**. While all variants of ACE2-Fc exhibited affinity to viral RBD protein, there were differences in their calculated EC_50_ values. Data are shown as the mean ± SD from triplicate experiments.

### 3.4. Competitive inhibition of pseudotyped viral transduction by R273A, H378A and E402A mutants of ΔACE2-Fc

ΔACE2-Fc Arg273Ala, His378Ale and Glu402Ala, and wild-type ACE2-Fc proteins were further tested for their antiviral potency. We conducted a series of viral inhibition assays using a pseudotyped reporter virus decorated with SARS-CoV-2 spike protein. The virus was able to transduce HEK293 cells that express full-length receptor ACE2 (See Methods). Each of the four ACE2-Fc variants in a range of concentrations was added to culture medium to test the potential of viral inhibition. As expected, all variants showed similar levels of efficacy to block pseudoviral transduction (**Fig4A**), with wild-type ACE2-Fc had a leading IC_50_ of 0.13 µg/mL, followed by His378Ala, Arg273Ala and Glu402Ala with their IC_50_s of 0.16 µg/mL, 0.19 µg/mL and 0.25 µg/mL, respectively (**Fig4B**).

**Figure 4.**
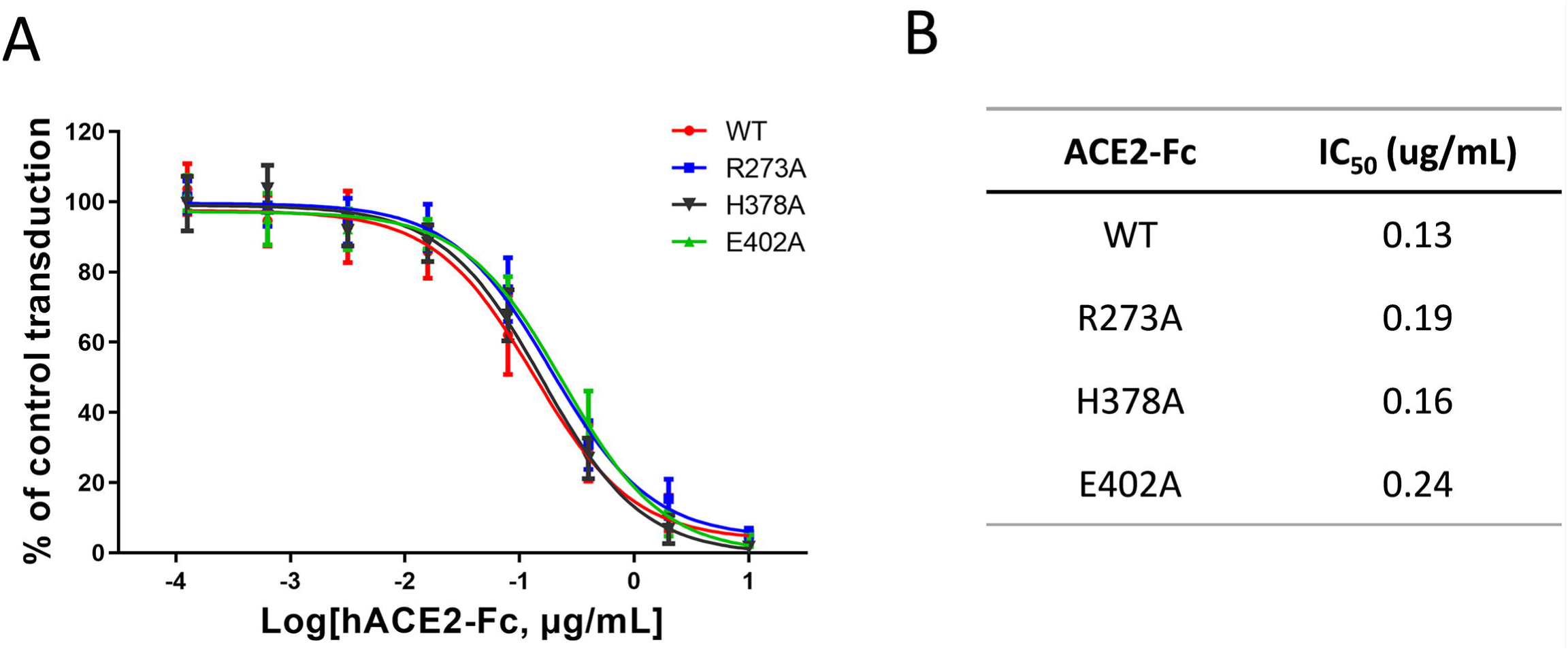
Inhibition of a pseudotyped virus by wild-type ACE2-Fc and ΔACE2-Fc variants. **A**. The transduction activity of a pseudotyped virus expressing SARS-CoV-2 spike protein to HEK293 cells expressing receptor ACE2 was measured through a firefly luciferase reporter. The cell transduction assays were performed in the presence of various concentrations of individual ACE2-Fc variants. **B**. IC_50_ values were calculated based on calculated ACE2-Fc concentrations needed to inhibit 50% reporter activity. Data are shown as the mean ± SD from triplicate experiments.

### 3.5. Pharmacokinetics of lead ΔACE2-Fc proteins

We have previously shown the in vivo longevity of mouse ACE2-Fc, as well as the fact that mouse FcRn recognizes human Fc[13]. Here we intravenously injected the ACE2-Fc variants in mice and measured pharmacokinetics of biologics. All three ΔACE2-Fc and their wild-type control exhibited long half-lives in the range between 52.61 hrs and 69.88 hrs (**Fig5**), consistent with the expectation for Fc-fusion proteins.

**Figure 5.**
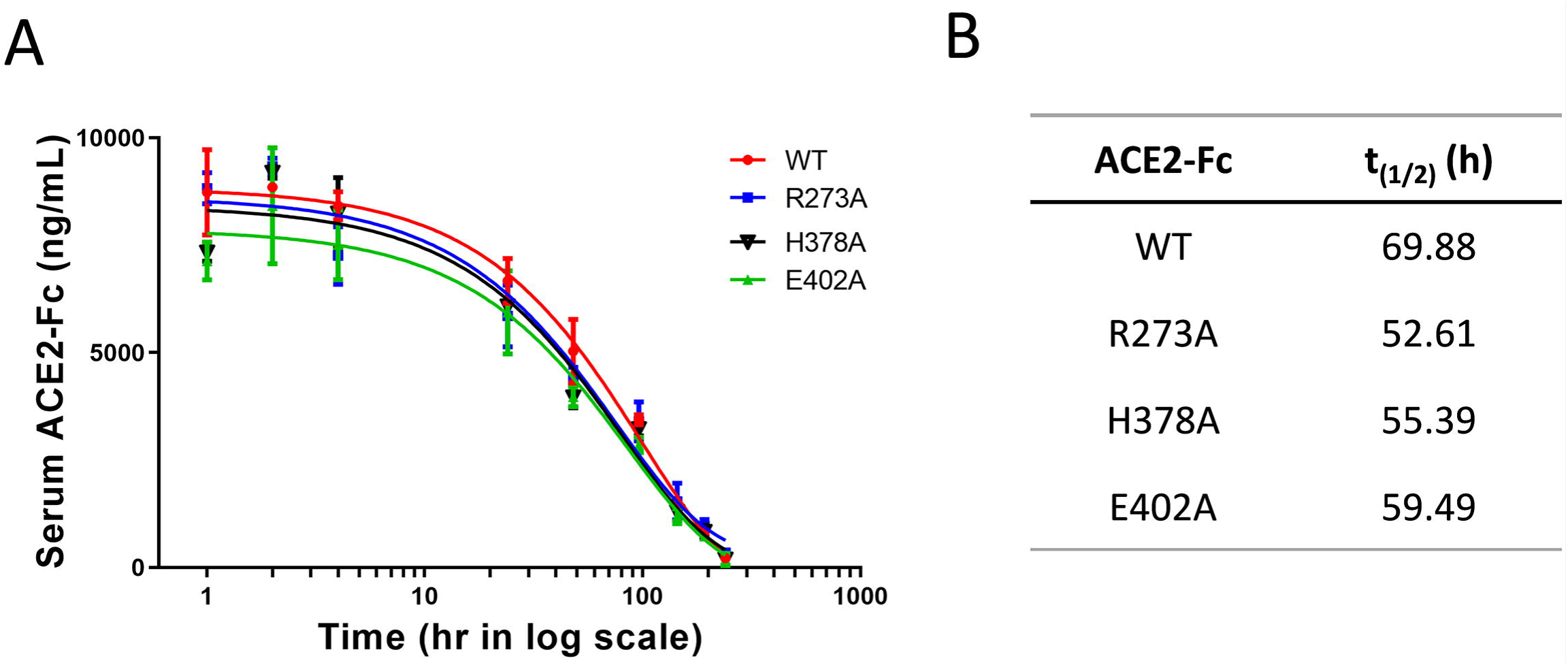
Pharmacokinetics of wild-type ACE2-Fc and ΔACE2-Fc variants. **A**. After a bolus *i.v.* injection of the listed ACE2-Fc variants in mice, drug concentration in blood was monitored over a period time. **B**. t_(1/2)_ values of individual biologics were calculated by GraphPad Prism software. Data are shown as the mean ± SD (n=3).

## 4. Discussions

Our study followed a design strategy of screening for ACE2-Fc variants of having an exclusive SARS-CoV-2 affinity with the absence of enzymatic activity towards vasoactive substrates. Based on the structure of ACE2’s catalytic center, we selected a total of 9 residues to be individually replaced with alanine. These included 3 residues with their side chains binding to divalent cation and 6 residues that line the substrate pocket. We used a surrogate fluorogenic substrate as well as two physiological substrates of ACE2 in reactions and discovered an unexpected substrate-dependent inactivation among individual mutants of ACE2-Fc. The screening identified three loss-of-activity variants (ΔACE2-Fc), one with a mutation of substrate-binding site and two others having impairment of cation-binding. All three lead candidates maintained their binding capacity towards SARS-CoV-2 spike protein and inhibited the transduction of a pseudotyped reporter virus.

Although we focused on inactivating ACE2 enzymatic activity to separate its actions on RAAS from SARS-CoV-2 neutralization, it has been widely speculated that the dual actions may benefit treatment of COVID-19. As ACE2 catalyzes the conversion of AngII to Ang-(1-7), therapeutic hrsACE2 or ACE2-Fc will change the balance from AngII-mediated stimulation of AT1 receptor to AT2 and/or Mas receptor activation, which may reduce pulmonary dysfunction due to AT1 associated inflammatory responses, lung edema and ARDS[15, 41-44]. Since there is no clinical data on ACE2-derived antiviral therapies, the hypotheses about beneficial RAAS inhibition are based on observations of COVID-19 patients who are on existing ACEI or ARB treatments. The general consensus is that these patients should continue RAAS blockade therapies for treating comorbidities during recovering from viral infection.

One of the main benefits of ACE2-Fc fusion construction is its long-acting time as compared to recombinant ACE2 without the tag[13]. It is expected to provide important assurance of sufficient drug levels to counteract the fluctuating levels of virions in patients, particularly during viremia. Based on clinical knowledge of Fc-tagged Factor VIII (ELOCTATE®) used in hemophilia A patients, dosing at 3-5 day intervals is sufficient to maintain a high blood level of the drug (US FDA recommendation).

The structural arrangement of ACE2-Fc resembles that of an antibody, with the replacement of antigen-binding Fab portion of antibody with ACE2 to bind SARS-CoV-2 spike. Meanwhile, the Fc portion can potentially induce immunological clearance of the virus, which, together in a fusion with ACE2, may be an effective immunoadhesin[11, 45] to trigger complement activation, antibody-mediated cytotoxicity and opsonization, and agglutination of targets. With respect to the potential antibody-like benefits of ACE2-Fc, we compare our overall strategy with existing CD4 immunoadhesin (termed PRO 542) that has existing clinical data for the treatment of HIV infection[46-50]. PRO 542 (CD4-IgG2/Fc with CD4 targeting HIV gp120) antiviral is a tetravalent fusion protein using the constant region of IgG2 as opposed to IgG1 of ACE2-Fc. One notable difference is that IgG2 has extremely low affinity to FcγRs on phagocytic cells, while both IgG1 and IgG2 can activate complement. With regard to our proof-of-principle study of ACE2-Fc as candidate antiviral drugs, it is important to point out that in terms of choices of the Fc tag, there are alternative strategies in recombinant construction. Nevertheless, the inclusion of immunoadhesin potentials of the antiviral may have caveats. While it may certainly boost immune clearance of the virus, such as through FcγR’s selective binding of clustered Fc, it may also elevate complement and cytokine responses to further aggravate the inflammation. Although these adverse side effects can be mitigated through modifications of the Fc domain, the ultimate therapeutic effects in the context of individual patients’ conditions can only be determined through rigorous clinical studies.

With regard to drug toxicity, our mouse study of repeated doses of ACE2-Fc in mice had showed the biologic to be well tolerated for up to two months[13]. However, we cannot extrapolate that its will also be safe COVID-19 patients. As we consider it is not a simple neutralizing agent of the virus, its bifurcated ACE2 head groups can possibly trigger agglutination of the virus that can potentially aggravate the hypercoagulable state, making the drug less tolerable in these conditions.

From the perspective of recombinant manufacturing, there are challenges ahead for the simple fact that ACE2-Fc is a large protein (∼130 kDa as a monomer and ∼260 kDa as a dimer). Furthermore, cocrystal structures of ACE2 with an inhibitor showed large movements of the two lobes as compared to the Apo structures[40], suggesting an intrinsic instability of ACE2 protein. Also of note is an earlier study by Lei et al using double mutations of His374 and His378 of the zinc-binding pocket for neutralization of SARS-CoV-2[6]. However, the ion is important in maintaining protein structure and stability of metallopeptidases[51] [52, 53]. This double mutation, as well as our His378Ala and Glu402Ala single mutations of the zinc-binding pocket may potentially suffer protein instability problems. On the other hand, Arg273Ala mutant that is predicted to change the substrate pocket will likely have a milder impact on overall protein stability, which is an important parameter in pharmaceutical production.

## Author statement

**Pan Liu**: Conceptualization, Methodology, Formal analysis, Writing – original draft. **Xinfang Xie**: Validation, Writing – review & editing. **Li Gao**: Validation, Writing – review & editing. **Jing Jin**: Conceptualization, Methodology, Project administration, Supervision, Writing – original draft.

## Acknowledgements

The design of the study was inspired by Dr. Robert Kruse’s F1000Research article. We thank Drs. Danial Batlle and Jan Wysocki for their insightful comments.

## Supplementary figures

**Supplementary Figure S1.**
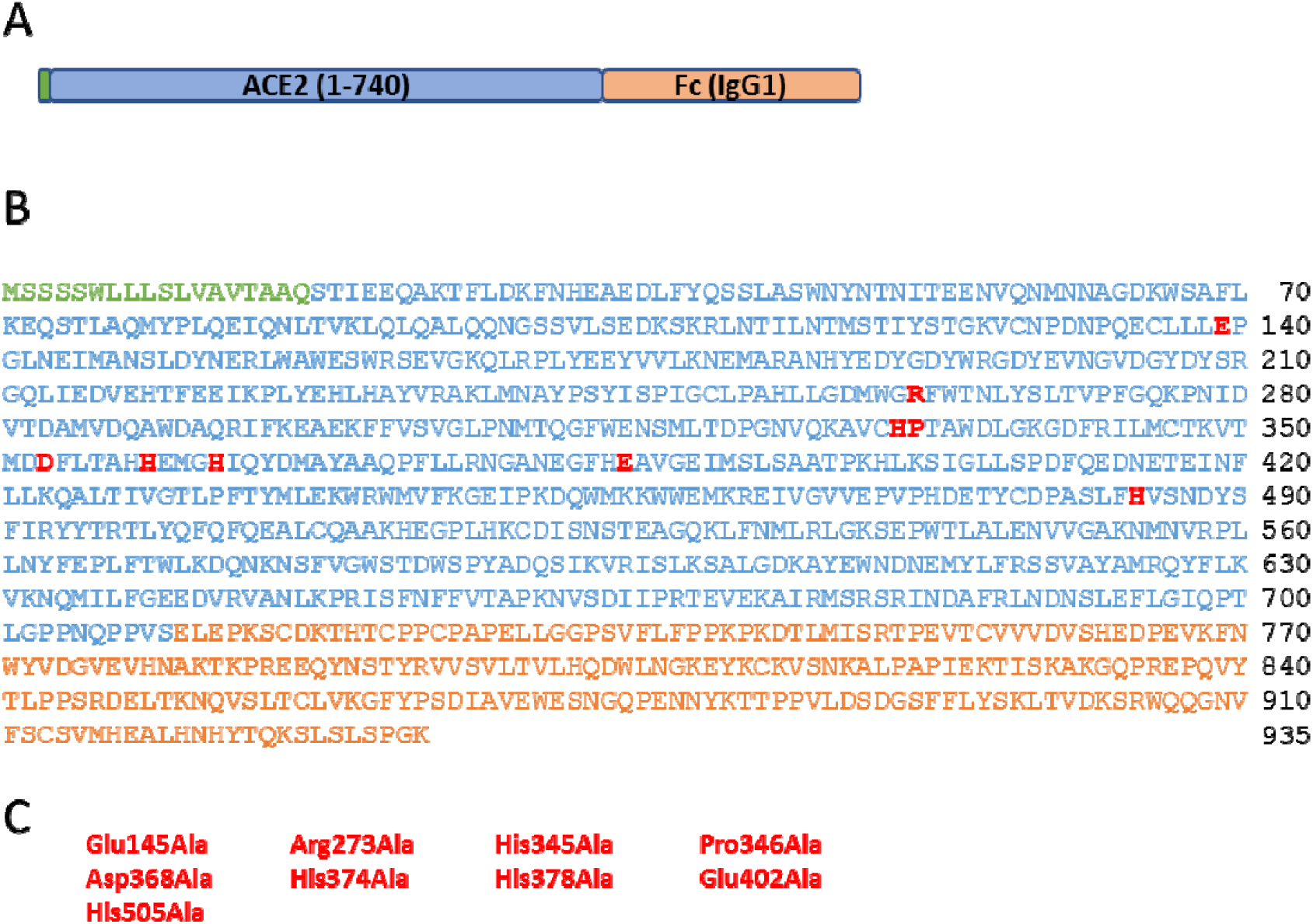
Summary of ACE2-Fc mutagenesis. **A**. Schematics of ACE2-Fc fusion construct: in an N-to-C-terminus order, ACE2-Fc is comprised of signal peptide and ACE2 ectodomain of human sequence, followed by Fc derived from human IgG1. **B**. The amino acid sequence of wild-type ACE2-Fc, showing signal peptide, ACE2 ectodomain and Fc with distinct font colors. The single amino acids in red fonts were individually mutated to alanine. C. List of the 9 mutants of ACE2-Fc.

**Supplementary Figure S2.**
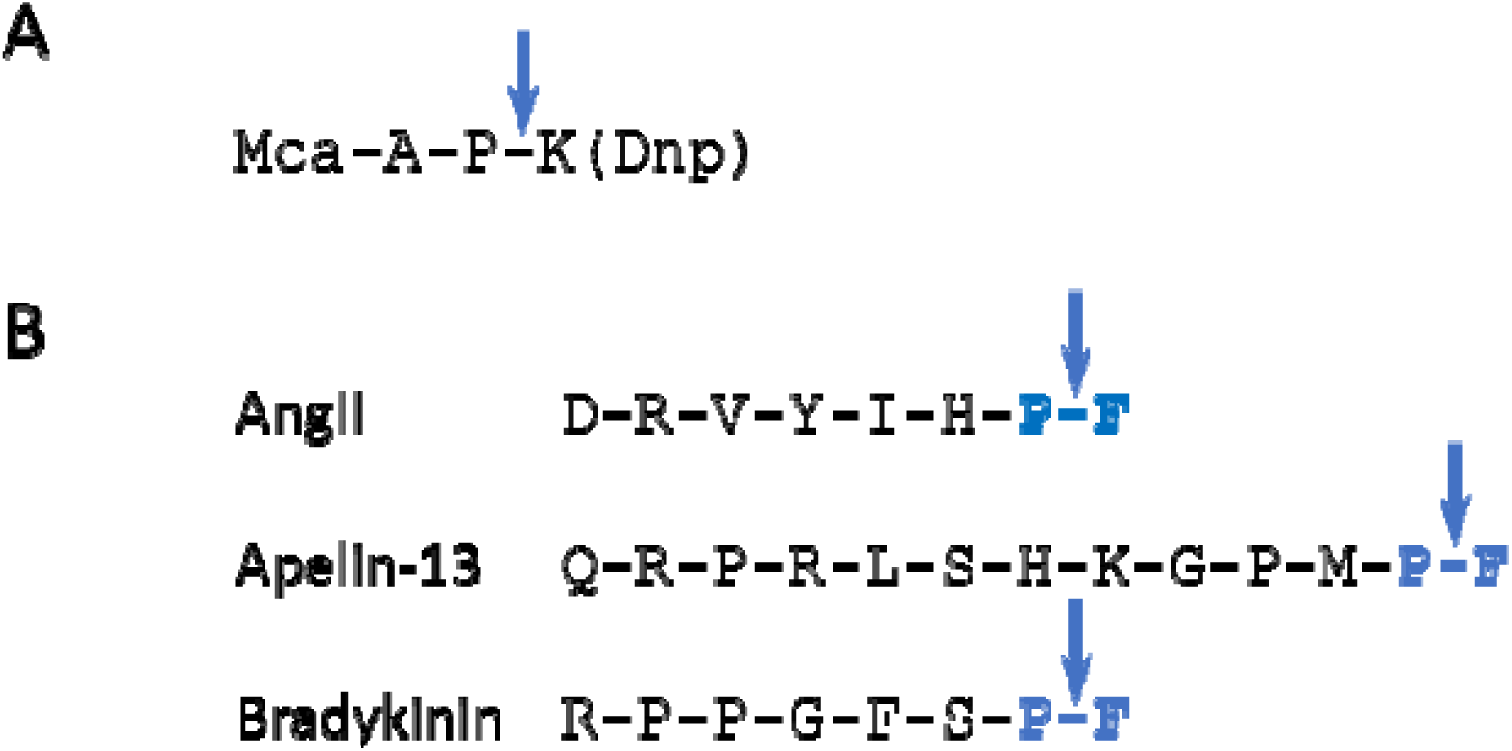
The consensus substrate motif of ACE2. **A**. ACE2 cleaves surrogate peptide of Mca-APK-(Dnp) between proline (P) and lysine (K) residues (arrow). **B**. ACE2 cleaves proline-phenylalanine (P-F) peptide bonds at the C-termini of its physiological peptides.

## References

[1] R. Yan, Y. Zhang, Y. Li, L. Xia, Y. Guo, Q. Zhou, Structural basis for the recognition of SARS-CoV-2 by full-length human ACE2, Science 367(6485) (2020) 1444–1448.

[2] D. Wrapp, N. Wang, K.S. Corbett, J.A. Goldsmith, C.L. Hsieh, O. Abiona, B.S. Graham, J.S. McLellan, Cryo-EM structure of the 2019-nCoV spike in the prefusion conformation, Science 367(6483) (2020) 1260–1263.

[3] P. Zhou, X.L. Yang, X.G. Wang, B. Hu, L. Zhang, W. Zhang, H.R. Si, Y. Zhu, B. Li, C.L. Huang, H.D. Chen, J. Chen, Y. Luo, H. Guo, R.D. Jiang, M.Q. Liu, Y. Chen, X.R. Shen, X. Wang, X.S. Zheng, K. Zhao, Q.J. Chen, F. Deng, L.L. Liu, B. Yan, F.X. Zhan, Y.Y. Wang, G.F. Xiao, Z.L. Shi, A pneumonia outbreak associated with a new coronavirus of probable bat origin, Nature 579(7798) (2020) 270–273.

[4] M. Hoffmann, H. Kleine-Weber, S. Schroeder, N. Kruger, T. Herrler, S. Erichsen, T.S. Schiergens, G. Herrler, N.H. Wu, A. Nitsche, M.A. Muller, C. Drosten, S. Pohlmann, SARS-CoV-2 Cell Entry Depends on ACE2 and TMPRSS2 and Is Blocked by a Clinically Proven Protease Inhibitor, Cell 181(2) (2020) 271–280 e8.

[5] V. Monteil, H. Kwon, P. Prado, A. Hagelkruys, R.A. Wimmer, M. Stahl, A. Leopoldi, E. Garreta, C. Hurtado Del Pozo, F. Prosper, J.P. Romero, G. Wirnsberger, H. Zhang, A.S. Slutsky, R. Conder, N. Montserrat, A. Mirazimi, J.M. Penninger, Inhibition of SARS-CoV-2 Infections in Engineered Human Tissues Using Clinical-Grade Soluble Human ACE2, Cell 181(4) (2020) 905–913 e7.

[6] C. Lei, K. Qian, T. Li, S. Zhang, W. Fu, M. Ding, S. Hu, Neutralization of SARS-CoV-2 spike pseudotyped virus by recombinant ACE2-Ig, Nat Commun 11(1) (2020) 2070.

[7] N. Iwanaga, L. Cooper, L. Rong, B. Beddingfield, J. Crabtree, R.A. Tripp, J.K. Kolls, Novel ACE2-IgG1 fusions with improved activity against SARS-CoV2, bioRxiv (2020).

[8] A. Khan, C. Benthin, B. Zeno, T.E. Albertson, J. Boyd, J.D. Christie, R. Hall, G. Poirier, J.J. Ronco, M. Tidswell, K. Hardes, W.M. Powley, T.J. Wright, S.K. Siederer, D.A. Fairman, D.A. Lipson, A.I. Bayliffe, A.L. Lazaar, A pilot clinical trial of recombinant human angiotensin-converting enzyme 2 in acute respiratory distress syndrome, Crit Care 21(1) (2017) 234.

[9] A.R. Hemnes, A. Rathinasabapathy, E.A. Austin, E.L. Brittain, E.J. Carrier, X. Chen, J.P. Fessel, C.D. Fike, P. Fong, N. Fortune, R.E. Gerszten, J.A. Johnson, M. Kaplowitz, J.H. Newman, R. Piana, M.E. Pugh, T.W. Rice, I.M. Robbins, L. Wheeler, C. Yu, J.E. Loyd, J. West, A potential therapeutic role for angiotensin-converting enzyme 2 in human pulmonary arterial hypertension, Eur Respir J 51(6) (2018).

[10] K.K. Chan, D. Dorosky, P. Sharma, S.A. Abbasi, J.M. Dye, D.M. Kranz, A.S. Herbert, E. Procko, Engineering human ACE2 to optimize binding to the spike protein of SARS coronavirus 2, Science (2020).

[11] R.L. Kruse, Therapeutic strategies in an outbreak scenario to treat the novel coronavirus originating in Wuhan, China, F1000Res 9 (2020) 72.

[12] X. Miao, Y. Luo, X. Huang, S.M.Y. Lee, Z. Yuan, Y. Tang, L. Chen, C. Wang, F. Wu, Y. Xu, W. Jiang, W. Gao, X. Song, Y. Yan, T. Pang, C. Chen, Y. Zou, W. Fu, L. Wan, J. Gilbert-Jaramillo, M. Knight, T.K. Tan, P. Rijal, A. Townsend, J. Sun, X. Liu, W. James, A. Tsun, Y. Xu, A novel biparatopic hybrid antibody-ACE2 fusion that blocks SARS-CoV-2 infection: implications for therapy, MAbs 12(1) (2020) 1804241.

[13] P. Liu, J. Wysocki, T. Souma, M. Ye, V. Ramirez, B. Zhou, L.D. Wilsbacher, S.E. Quaggin, D. Batlle, J. Jin, Novel ACE2-Fc chimeric fusion provides long-lasting hypertension control and organ protection in mouse models of systemic renin angiotensin system activation, Kidney Int 94(1) (2018) 114–125.

[14] Y. Imai, K. Kuba, S. Rao, Y. Huan, F. Guo, B. Guan, P. Yang, R. Sarao, T. Wada, H. Leong-Poi, M.A. Crackower, A. Fukamizu, C.C. Hui, L. Hein, S. Uhlig, A.S. Slutsky, C. Jiang, J.M. Penninger, Angiotensin-converting enzyme 2 protects from severe acute lung failure, Nature 436(7047) (2005) 112–6.

[15] G.P. Rossi, V. Sanga, M. Barton, Potential harmful effects of discontinuing ACE-inhibitors and ARBs in COVID-19 patients, Elife 9 (2020).

[16] T.J. Guzik, S.A. Mohiddin, A. Dimarco, V. Patel, K. Savvatis, F.M. Marelli-Berg, M.S. Madhur, M. Tomaszewski, P. Maffia, F. D’Acquisto, S.A. Nicklin, A.J. Marian, R. Nosalski, E.C. Murray, B. Guzik, C. Berry, R.M. Touyz, R. Kreutz, D.W. Wang, D. Bhella, O. Sagliocco, F. Crea, E.C. Thomson, I.B. McInnes, COVID-19 and the cardiovascular system: implications for risk assessment, diagnosis, and treatment options, Cardiovasc Res (2020).

[17] E.R. Lumbers, S.J. Delforce, K.G. Pringle, G.R. Smith, The Lung, the Heart, the Novel Coronavirus, and the Renin-Angiotensin System; The Need for Clinical Trials, Front Med (Lausanne) 7 (2020) 248.

[18] A. Cai, B. McClafferty, J. Benson, D. Ramgobin, R. Kalayanamitra, Z. Shahid, A. Groff, C.S. Aggarwal, R. Patel, H. Polimera, R. Vunnam, R. Golamari, N. Sahu, D. Bhatt, R. Jain, COVID-19: Catastrophic Cause of Acute Lung Injury, S D Med 73(6) (2020) 252–260.

[19] Y.Y. Zheng, Y.T. Ma, J.Y. Zhang, X. Xie, COVID-19 and the cardiovascular system, Nat Rev Cardiol 17(5) (2020) 259–260.

[20] A.H.J. Danser, M. Epstein, D. Batlle, Renin-Angiotensin System Blockers and the COVID-19 Pandemic: At Present There Is No Evidence to Abandon Renin-Angiotensin System Blockers, Hypertension 75(6) (2020) 1382–1385.

[21] M.A. Sparks, A. South, P. Welling, J.M. Luther, J. Cohen, J.B. Byrd, L.M. Burrell, D. Batlle, L. Tomlinson, V. Bhalla, M.N. Rheault, M.J. Soler, S. Swaminathan, S. Hiremath, Sound Science before Quick Judgement Regarding RAS Blockade in COVID-19, Clin J Am Soc Nephrol 15(5) (2020) 714–716.

[22] C. Bavishi, T.M. Maddox, F.H. Messerli, Coronavirus Disease 2019 (COVID-19) Infection and Renin Angiotensin System Blockers, JAMA Cardiol (2020).

[23] M. Vaduganathan, O. Vardeny, T. Michel, J.J.V. McMurray, M.A. Pfeffer, S.D. Solomon, Renin-Angiotensin-Aldosterone System Inhibitors in Patients with Covid-19, N Engl J Med 382(17) (2020) 1653–1659.

[24] A.M. South, T.M. Brady, J.T. Flynn, ACE2 (Angiotensin-Converting Enzyme 2), COVID-19, and ACE Inhibitor and Ang II (Angiotensin II) Receptor Blocker Use During the Pandemic: The Pediatric Perspective, Hypertension 76(1) (2020) 16–22.

[25] M.R. Mehra, S.S. Desai, S. Kuy, T.D. Henry, A.N. Patel, Cardiovascular Disease, Drug Therapy, and Mortality in Covid-19, N Engl J Med 382(25) (2020) e102.

[26] A.M. South, D.I. Diz, M.C. Chappell, COVID-19, ACE2, and the cardiovascular consequences, Am J Physiol Heart Circ Physiol 318(5) (2020) H1084–H1090.

[27] R. Sommerstein, M.M. Kochen, F.H. Messerli, C. Grani, Coronavirus Disease 2019 (COVID-19): Do Angiotensin-Converting Enzyme Inhibitors/Angiotensin Receptor Blockers Have a Biphasic Effect?, J Am Heart Assoc 9(7) (2020) e016509.

[28] F. Zhou, T. Yu, R. Du, G. Fan, Y. Liu, Z. Liu, J. Xiang, Y. Wang, B. Song, X. Gu, L. Guan, Y. Wei, H. Li, X. Wu, J. Xu, S. Tu, Y. Zhang, H. Chen, B. Cao, Clinical course and risk factors for mortality of adult inpatients with COVID-19 in Wuhan, China: a retrospective cohort study, Lancet 395(10229) (2020) 1054–1062.

[29] J. Guo, Z. Huang, L. Lin, J. Lv, Coronavirus Disease 2019 (COVID-19) and Cardiovascular Disease: A Viewpoint on the Potential Influence of Angiotensin-Converting Enzyme Inhibitors/Angiotensin Receptor Blockers on Onset and Severity of Severe Acute Respiratory Syndrome Coronavirus 2 Infection, J Am Heart Assoc 9(7) (2020) e016219.

[30] M. Noris, A. Benigni, G. Remuzzi, The case of complement activation in COVID-19 multiorgan impact, Kidney Int (2020).

[31] K. Renu, P.L. Prasanna, A. Valsala Gopalakrishnan, Coronaviruses pathogenesis, comorbidities and multi-organ damage – A review, Life Sci 255 (2020) 117839.

[32] S. Zaim, J.H. Chong, V. Sankaranarayanan, A. Harky, COVID-19 and Multiorgan Response, Curr Probl Cardiol 45(8) (2020) 100618.

[33] H. Bosmuller, S. Traxler, M. Bitzer, H. Haberle, W. Raiser, D. Nann, L. Frauenfeld, A. Vogelsberg, K. Klingel, F. Fend, The evolution of pulmonary pathology in fatal COVID-19 disease: an autopsy study with clinical correlation, Virchows Arch (2020).

[34] T. Menter, J.D. Haslbauer, R. Nienhold, S. Savic, H. Hopfer, N. Deigendesch, S. Frank, D. Turek, N. Willi, H. Pargger, S. Bassetti, J.D. Leuppi, G. Cathomas, M. Tolnay, K.D. Mertz, A. Tzankov, Post-mortem examination of COVID19 patients reveals diffuse alveolar damage with severe capillary congestion and variegated findings of lungs and other organs suggesting vascular dysfunction, Histopathology (2020).

[35] P. Liu, J. Wysocki, P. Serfozo, M. Ye, T. Souma, D. Batlle, J. Jin, A Fluorometric Method of Measuring Carboxypeptidase Activities for Angiotensin II and Apelin-13, Sci Rep 7 (2017) 45473.

[36] A.C. Walls, Y.J. Park, M.A. Tortorici, A. Wall, A.T. McGuire, D. Veesler, Structure, Function, and Antigenicity of the SARS-CoV-2 Spike Glycoprotein, Cell 181(2) (2020) 281–292 e6.

[37] J. Shang, G. Ye, K. Shi, Y. Wan, C. Luo, H. Aihara, Q. Geng, A. Auerbach, F. Li, Structural basis of receptor recognition by SARS-CoV-2, Nature 581(7807) (2020) 221–224.

[38] J. Lan, J. Ge, J. Yu, S. Shan, H. Zhou, S. Fan, Q. Zhang, X. Shi, Q. Wang, L. Zhang, X. Wang, Structure of the SARS-CoV-2 spike receptor-binding domain bound to the ACE2 receptor, Nature 581(7807) (2020) 215–220.

[39] Q. Wang, Y. Zhang, L. Wu, S. Niu, C. Song, Z. Zhang, G. Lu, C. Qiao, Y. Hu, K.Y. Yuen, Q. Wang, H. Zhou, J. Yan, J. Qi, Structural and Functional Basis of SARS-CoV-2 Entry by Using Human ACE2, Cell 181(4) (2020) 894–904 e9.

[40] P. Towler, B. Staker, S.G. Prasad, S. Menon, J. Tang, T. Parsons, D. Ryan, M. Fisher, D. Williams, N.A. Dales, M.A. Patane, M.W. Pantoliano, ACE2 X-ray structures reveal a large hinge-bending motion important for inhibitor binding and catalysis, J Biol Chem 279(17) (2004) 17996–8007.

[41] P. Namsolleck, G.N. Moll, Does activation of the protective Renin-Angiotensin System have therapeutic potential in COVID-19?, Mol Med 26(1) (2020) 80.

[42] M.K. Chung, S. Karnik, J. Saef, C. Bergmann, J. Barnard, M.M. Lederman, J. Tilton, F. Cheng, C.V. Harding, J.B. Young, N. Mehta, S.J. Cameron, K.R. McCrae, A.H. Schmaier, J.D. Smith, A. Kalra, S.K. Gebreselassie, G. Thomas, E.S. Hawkins, L.G. Svensson, SARS-CoV-2 and ACE2: The biology and clinical data settling the ARB and ACEI controversy, EBioMedicine 58 (2020) 102907.

[43] R. Sarzani, F. Giulietti, C. Di Pentima, P. Giordano, F. Spannella, Disequilibrium between the classic renin-angiotensin system and its opposing arm in SARS-CoV-2-related lung injury, Am J Physiol Lung Cell Mol Physiol 319(2) (2020) L325–L336.

[44] R.D. Lopes, A.V.S. Macedo, E.S.P.G.M. de Barros, R.J. Moll-Bernardes, A. Feldman, G. D’Andrea Saba Arruda, A.S. de Souza, D.C. de Albuquerque, L. Mazza, M.F. Santos, N.Z. Salvador, C.M. Gibson, C.B. Granger, J.H. Alexander, O.F. de Souza, B.C. investigators, Continuing versus suspending angiotensin-converting enzyme inhibitors and angiotensin receptor blockers: Impact on adverse outcomes in hospitalized patients with severe acute respiratory syndrome coronavirus 2 (SARS-CoV-2)--The BRACE CORONA Trial, Am Heart J 226 (2020) 49–59.

[45] A. Ashkenazi, S.M. Chamow, Immunoadhesins as research tools and therapeutic agents, Curr Opin Immunol 9(2) (1997) 195–200.

[46] S.M. Chamow, A.M. Duliege, A. Ammann, J.O. Kahn, J.D. Allen, J.W. Eichberg, R.A. Byrn, D.J. Capon, R.H. Ward, A. Ashkenazi, CD4 immunoadhesins in anti-HIV therapy: new developments, Int J Cancer Suppl 7 (1992) 69–72.

[47] C.V. Fletcher, J.G. DeVille, P.M. Samson, J.H. Moye, Jr., J.A. Church, H.M. Spiegel, P. Palumbo, T. Fenton, M.E. Smith, B. Graham, J.M. Kraimer, W.T. Shearer, P.S.G. Pediatric Aids Clinical Trials Group, Nonlinear pharmacokinetics of high-dose recombinant fusion protein CD4-IgG2 (PRO 542) observed in HIV-1-infected children, J Allergy Clin Immunol 119(3) (2007) 747–50.

[48] J.M. Jacobson, R.J. Israel, I. Lowy, N.A. Ostrow, L.S. Vassilatos, M. Barish, D.N. Tran, B.M. Sullivan, T.J. Ketas, T.J. O’Neill, K.A. Nagashima, W. Huang, C.J. Petropoulos, J.P. Moore, P.J. Maddon, W.C. Olson, Treatment of advanced human immunodeficiency virus type 1 disease with the viral entry inhibitor PRO 542, Antimicrob Agents Chemother 48(2) (2004) 423–9.

[49] J.M. Jacobson, I. Lowy, C.V. Fletcher, T.J. O’Neill, D.N. Tran, T.J. Ketas, A. Trkola, M.E. Klotman, P.J. Maddon, W.C. Olson, R.J. Israel, Single-dose safety, pharmacology, and antiviral activity of the human immunodeficiency virus (HIV) type 1 entry inhibitor PRO 542 in HIV-infected adults, J Infect Dis 182(1) (2000) 326–9.

[50] W.T. Shearer, R.J. Israel, S. Starr, C.V. Fletcher, D. Wara, M. Rathore, J. Church, J. DeVille, T. Fenton, B. Graham, P. Samson, S. Staprans, J. McNamara, J. Moye, P.J. Maddon, W.C. Olson, Recombinant CD4-IgG2 in human immunodeficiency virus type 1-infected children: phase 1/2 study. The Pediatric AIDS Clinical Trials Group Protocol 351 Study Team, J Infect Dis 182(6) (2000) 1774–9.

[51] S. Fridrich, K. Karmilin, W. Stocker, Handling Metalloproteinases, Curr Protoc Protein Sci 83 (2016) 21 16 1–21 16 20.

[52] F. Namuswe, J.M. Berg, Secondary interactions involving zinc-bound ligands: roles in structural stabilization and macromolecular interactions, J Inorg Biochem 111 (2012) 146–9.

[53] K.A. McCall, C. Huang, C.A. Fierke, Function and mechanism of zinc metalloenzymes, J Nutr 130(5S Suppl) (2000) 1437S–46S.

