## Supplemental Figures for "Designed Variants of ACE2-Fc that Decouple Anti-SARS-CoV-2 Activities from Unwanted Cardiovascular Effects"

A

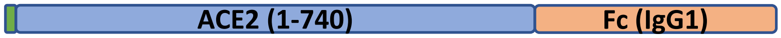


B

MSSSSWLLLSLVAVTAAQSTIEEQAKTFLDKFNHEAEDLFYQSSLASWNYNTNITEENVQNMNNAGDKWSAFL 70

KEQSTLAQMYPLQEIQNLTVKLQLQALQQNGSSVLSEDKSKRLNTILNTMSTIYSTGKVCNPDNPQECLLL**E**P 140

GLNEIMANSLDYNERLWAWESWRSEVGKQLRPLYEEYVVLKNEMARANHYEDYGDYWRGDYEVNGVDGYDYSR 210

GQLIEDVEHTFEEIKPLYEHLHAYVRAKLMNAYPSYISPIGCLPAHLLGDMWG**R**FWTNLYSLTVPFGQKPNID 280

VTDAMVDQAWDAQRIFKEAEKFFVSVGLPNMTQGFWENSMLTDPGNVQKAVC**HP**TAWDLGKGDFRILMCTKVT 350

MD**D**FLTAH**H**EMG**H**IQYDMAYAAQPFLLRNGANEGFH**E**AVGEIMSLSAATPKHLKSIGLLSPDFQEDNETEINF 420

LLKQALTIVGTLPFTYMLEKWRWMVFKGEIPKDQWMKKWWEMKREIVGVVEPVPHDETYCDPASLF**H**VSNDYS 490

FIRYYTRTLYQFQFQEALCQAAKHEGPLHKCDISNSTEAGQKLFNMLRLGKSEPWTLALENVVGAKNMNVRPL 560

LNYFEPLFTWLKDQNKNSFVGWSTDWSPYADQSIKVRISLKSALGDKAYEWNDNEMYLFRSSVAYAMRQYFLK 630

VKNQMILFGEEDVRVANLKPRISFNFFVTAPKNVSDIIPRTEVEKAIRMSRSRINDAFRLNDNSLEFLGIQPT 700

LGPPNQPPVSELEPKSCDKTHTCPPCPAPELLGGPSVFLFPPKPKDTLMISRTPEVTCVVVDVSHEDPEVKFN 770

WYVDGVEVHNAKTKPREEQYNSTYRVVSVLTVLHQDWLNGKEYKCKVSNKALPAPIEKTISKAKGQPREPQVY 840

TLPPSRDELTKNQVSLTCLVKGFYPSDIAVEWESNGQPENNYKTTPPVLDSDGSFFLYSKLTVDKSRWQQGNV 910

FSCSVMHEALHNHYTQKSLSLSPGK 935

C

**Glu145Ala** **Arg273Ala** **His345Ala** **Pro346Ala**

**Asp368Ala** **His374Ala** **His378Ala** **Glu402Ala**

**His505Ala**

**Supplementary Figure S1. Summary of ACE2-Fc mutagenesis. A**. Schematics of ACE2-Fc fusion construct: in an N-to-C-terminus order, ACE2-Fc is comprised of signal peptide and ACE2 ectodomain of human sequence, followed by Fc derived from human IgG1. **B**. The amino acid sequence of wild-type ACE2-Fc, showing signal peptide, ACE2 ectodomain and Fc with distinct font colors. The single amino acids in red fonts were individually mutated to alanine. **C**. List of the 9 mutants of ACE2-Fc.

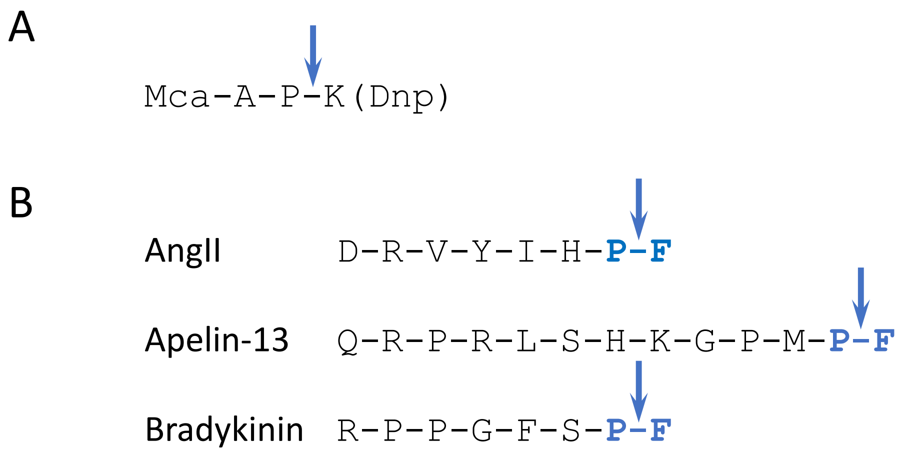


**Supplementary Figure S2. The consensus substrate motif of ACE2 . A**. ACE2 cleaves surrogate peptide of Mca-APK-(Dnp) between proline (P) and lysine (K) residues (arrow). **B**. ACE2 cleaves proline-phenylalanine (P-F) peptide bonds at the C-termini of its physiological peptides.
